# Monitoring macrophage polarization with gene expression reporters and bioluminescence phasor analysis

**DOI:** 10.1101/2024.06.10.598305

**Authors:** Giulia Tedeschi, Mariana X. Navarro, Lorenzo Scipioni, Tanvi K. Sondhi, Jennifer A. Prescher, Michelle A. Digman

**Affiliations:** Laboratory for Fluorescence Dynamics, Biomedical Engineering Department, University of California, Irvine, Irvine, CA 92617 (USA); Department of Chemistry, University of California, Irvine, Irvine, CA 92617 (USA); Department of Molecular Biology and Biochemistry, University of California, Irvine, Irvine, CA 92617 (USA); Department of Pharmaceutical Sciences, University of California, Irvine, Irvine, CA 92617

**Author notes:** Equal contribution (Giulia Tedeschi, Mariana X. Navarro).

**Keywords:** Bioluminescence, Single-cell, Microscopy, Phasor, Macrophages, Gene Reporters

## Abstract

Macrophages exhibit a spectrum of behaviors upon activation and are generally classified as one of two types: inflammatory (M1) or anti-inflammatory (M2). Tracking these phenotypes in living cells can provide insight into immune function, but remains a challenging pursuit. Existing methods are mostly limited to static readouts or difficult to employ for multiplexed imaging in complex 3D environments while maintaining cellular resolution. We aimed to fill this void using bioluminescent technologies. Here we report genetically engineered luciferase reporters for long-term monitoring of macrophage polarization via spectral phasor analysis. M1- and M2-specific promoters were used to drive the expression of bioluminescent enzymes in macrophage cell lines. The readouts were multiplexed and discernable in both 2D and 3D formats with single cell resolution in living samples. Collectively, this work expands the toolbox of methods for monitoring macrophage polarization and provides a blueprint for monitoring other multifaceted networks in heterogeneous environments.

## Introduction

Macrophages are intricately involved in immunity and inflammation,^1–4^ tissue development, ^5–7^ wound repair,^8^ and homeostasis.^9,10^ These cells have a wide range of functions that are classified as either pro-inflammatory (M1) or anti-inflammatory (M2). M1 macrophages are typically associated with pathogen killing, while M2 macrophages play central roles in tissue healing and growth. Activation of macrophages into M1 and M2 phenotypes is induced by external stimuli. Among the most well-known is lipopolysaccharide (LPS), an inflammatory stimulus that polarizes macrophages to an M1 state by triggering the macrophages to secrete different cytokines.^11,12^ By contrast, macrophage exposure to IL-4/IL-13 cytokines drives M2 polarization.^13^ Both of these cytokines bind to receptors on the cell surface and induce a variety of signaling pathways, dampening the inflammatory response.^14^

Polarized macrophages have a broad range of functions in distinct environments, including wounded tissue and tumors.^15^ In these settings, M2 macrophages often play key roles in the final stages of tissue remodeling and resolution of inflammation.^15,16^ M2 polarized macrophages can thus influence tumor proliferation and immune evasion. M1 macrophages, by contrast, tend to produce inflammatory factors and thus exhibit more anti-tumor effects.^17,18^ M1/M2 polarization occurs along a spectrum, though, and macrophages can lie anywhere between a completely M1 or M2 phenotype. Being able to track precise polarization statuses—in real-time—could provide critical information on cell state in a tumor microenvironment and inform on therapeutic options.^19^

While macrophage plasticity allows for specific responses to environmental stimuli, visualizing these processes over extended periods and directly in live tissue environments remains limited. In clinical practice, monitoring macrophage phenotypes is often based on static readouts such as flow cytometry or immunohistochemistry. Flow cytometry enables standardized quantification of macrophage phenotypes, but lacks spatial analysis. In contrast, immunohistochemistry (IHC) methods contain spatial information, but remains to be standardized, resulting in unreliable results across samples and researchers.^20^

Real-time imaging of macrophage function is possible using reporter genes and/or fluorescent dye labeling.^21^ Cell-specific promoters driving detectable gene expression have enabled long term tracking.^22^ Similarly, fluorescent dyes have been used for quantifying biomarkers and macrophage activities.^21,23^ Both approaches relay on external light sources for signal production, which can result in photobleaching and phototoxicity. Fluorescent reporters can also be difficult to apply in tissues owing to autofluorescence.^21^ Macrophage imaging is possible with more tissue-penetrant imaging modalities (e.g., PET and MRI), but these methods are limited in multiplexing and long term imaging capabilities.^21,24^

To address the need for improved visualization of macrophage behavior in heterogeneous environments, we leveraged bioluminescence imaging (BLI). Bioluminescence involves light production from the interaction between luciferase enzymes and luciferin small molecules^25^. No excitation light is required, making this technique well suited for imaging in tissue and other opaque environments. Several bioluminescence-based macrophage reporter cells have been developed for monitoring gene expression.^26–30^ However, most are limited to either *in vitro* model systems, tracking a single marker over time, or lack single cell resolution.^21,31^ In the absence of tools to monitor live cells and complex interactions in real time, there is often a disconnection between *in vitro* biomarkers and model systems and *in vivo* analysis of macrophage polarization.^21,32,33^

Here we report a strategy for live-cell imaging of macrophages using bioluminescence resonance energy transfer (BRET) and spectral phasor analysis. We developed two distinct BRET reporters that correlate with *NOS2* (M1) and *STAT6* (M2) expression. We demonstrated that the reporters can provide a readout on macrophage polarization, using spectral phasor analysis to provide single cell readouts. The polarization status of the cell lines was confirmed by monitoring organelle features with fluorescence microscopy. We further demonstrated that the reporters and spectral phasor analysis could be used to examine macrophage status in a 3D cell culture model. Overall, our work provides a platform for multiplexed monitoring of immune cell polarization in a range of environments.

## Material and Methods

### General cloning methods

Promoter regions and genes of interest were amplified using polymerase chain reaction (PCR). The *STAT6* promoter region was amplified from the p4xSTAT6-Luc2P plasmid (Addgene #35554). The *NOS2* promoter region was amplified from the pGL2-NOS2Promoter-Luciferase plasmid (Addgene #19296). YeNL and CeNL were amplified from plasmids as previously described.^34^ Primer melting temperatures were calculated using a melting temperature (T_m_) calculator offered by New England BioLabs (https://tmcalculator.neb.com). All PCR reactions were performed using a BioRad C3000 Thermocycler using the following conditions: 1x Q5 Hot start DNA polymerase reaction buffer, dNTPs (0.8 mM) and Q5 Hot start DNA polymerase (1 U) in a total reaction volume of 50 μL, unless otherwise stated. The following thermal cycling conditions were used to amplify all inserts: 20 cycles of denaturation (95 °C, 30 s), annealing (60 °C over 30 s) and extension (72 °C, 180 s). The PCR products were purified via gel electrophoresis using 1% agarose gels and products were identified using GelRed® Nucleic Acid Gel Stain (Fisher Scientific).

The inserts were assembled into a vector for viral transduction (pLenti, Addgene #73582). Plasmids were digested with *Cla*I (New England Biolabs) and *Bam*HI (New England Biolabs) for 3 h at 37°C. The products were purified from remaining circular template via gel electrophoresis in 1% agarose gels. Inserts were assembled with linearized vectors using Gibson assembly.^35^ Gibson assembly master mixes were prepared following the recipe from Prather and coworkers (http://www.openwetware.org/wiki/Gibson_Assembly), with all materials purchased from New England BioLabs. For the assembly, 50 ng of linearized vector was combined with insert (2:1 insert:vector ratio) and added to 10 μL of master mix. The mixtures were incubated at 50°C for 1 h, then transformed. Ligated plasmids were transformed into the TOP10 strain of *E. coli* using the heat shock method. Colonies containing genes of interest were expanded overnight in 5 mL of LB broth supplemented with ampicillin (100 μg/mL). DNA was extracted using a Zymo Research Plasmid Miniprep kit and concentrations were measured with a Nanodrop 2000c spectrophotometer (Thermo Scientific). Sequencing analyses were used to confirm successful plasmid generation.

**Table 1.**
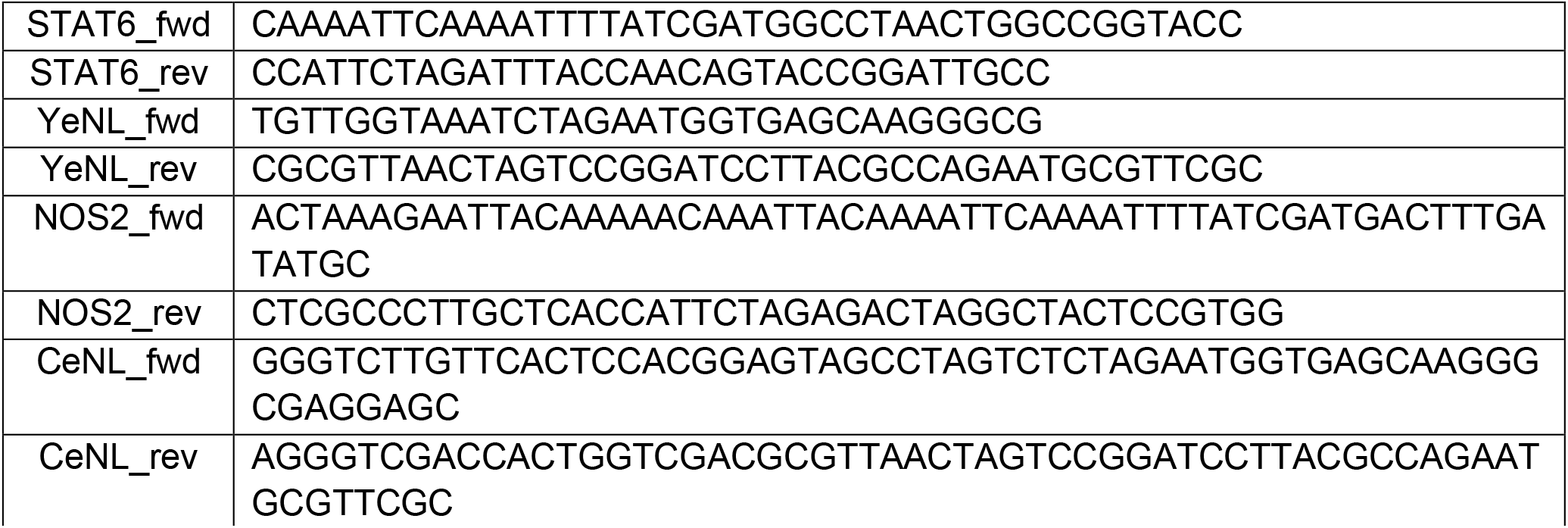
Primers used for amplification of the promoter regions and gene inserts. All primers were purchased from Integrated DNA Technologies, Inc. (San Diego, CA) and are written in the 5′ ⟶3′ direction.

### Mammalian cell culture

RAW264.7 (ATCC) were cultured in DMEM (Corning) supplemented with 10% (v/v) fetal bovine serum (FBS, Life Technologies), penicillin (100 U/mL) and streptomycin (100 μg/mL). Cells were maintained in a 5% (v/v) CO_2_ water-saturated incubator at 37 °C. RAW264.7 cells were serially passaged using Hank’s Balanced Salt Solution (HBSS, Gibco) and cell scrapers (Fisher Scientific). Cells were counted using an automated cell counter (Countess II, Invitrogen).

Cells were seeded (4 × 10^5^) in 6-well dishes (Corning). STAT6 and NOS2 plasmids were co-transfected with psPAX2 and pMD2.G with Lipofectamine 3000 (ThermoFisher) according to manufacturer’s instructions. About 16 h post transfection, the cell culture media was replaced with media to induce viral transduction (DMEM supplemented with 10 mM sodium butyrate, 20 mM HEPES, and 2 mM L-glutamine). After a 24–48 h incubation, the media was replaced with standard DMEM culture media, which was subsequently collected 24–48 h later. The media was spun down to pellet cell debris, then the supernatant containing the virus was stored at –80 °C or used immediately for transduction. Transduced cells were selected using puromycin (2–20 μg/mL).

### Macrophage stimulation and sample preparation for imaging

RAW264.7 cells were seeded in either a 12-well plate (Corning) or 8-well chambered coverglass (Ibidi). To stimulate to an M1 phenotype, lipopolysaccharide was added (LPS, 5 μg/mL, Invitrogen) and to an M2 phenotype, IL-4 (20 ng/mL, R&D Systems) and IL-13 (20 ng/mL, R&D Systems). After 18–24 h incubation, unless otherwise described, cells were either imaged on the TIRF microscope, as described below, or transferred to a 96-well plate for luminescence analysis. For the experiments with a mixture of cell populations, reporter cell types were mixed before plating in an 8-well slide. Final cell counts per well consisted of 1 × 10^5^ of each cell type (1:1 mixtures). Cell mixtures were then stimulated to an M1 or M2 phenotype as previously described.

### Embedding cells in collagen matrix

Collagen matrix was prepared from High Concentration Rat Tail I collagen (Corning) to be at a final concentration of 2.0 mg/mL (pH 7.4). For each condition, cells (3 ×10^5^ cells total) were washed with PBS (1X, Gibco), pelleted at 180 x*g*, resuspended in 150 μL of collagen solution per well and plated in 8-well chambers (Cellvis). The samples were kept at room temperature for 30 min and then incubated (37 °C, 5% v/v CO_2_) for 1 h to allow collagen polymerization. Media (200 μL) was then added on top of the collagen matrix and kept in the incubator until the moment of use.

### Luminometer bioluminescence imaging

Bioluminescence scans were performed using a Tecan Spark multimode microplate reader. Cells were plated in black 96-well plates (Grenier Bio One) and furimazine (Promega, Nano-Glo Luciferase Assay System) was added to each well. Immediately after administration of the luciferin, the plate was shaken (5 s) and luminescence scans were taken. Samples were analyzed in triplicate and data were exported to Microsoft Excel or Prism (GraphPad) for further analysis.

### Bioluminescence microscopy imaging

Bioluminescence phasor imaging was performed on an Olympus IX83 Total internal reflection fluorescence (TIRF) microscope equipped with two Optosplit II (Cairn) image splitters used in widefield mode. All components and imaging software are described in previous work^34^. Transduced RAW 264.7 reporter cells (2 × 10^5^ cells/well) were plated on an 8-well chambered cover glass (Ibidi). Medium was removed and replaced with a fresh stock of medium (300 μL) containing furimazine (25-50 μM). Five minutes post-media exchange, the cells were imaged with an Olympus TIRF microscope in widefield mode using a 20X or 10X air objective (Olympus UPlanSAPO 20X/0.75, Olympus UPlanSApo 10X/0.40) with further 2X magnification. All images were recorded with 10 s integration time and 20 frames total were collected per sample. Images were exported as TIFF files and analyzed as described below.

### Image processing and statistical analysis

Images exported in TIFF format and processed with a custom Python algorithm written in Google Colab with a workflow similar to that previously described^34^. Images were split from a single TIFF file into four channels corresponding to the sine- and cosine-filtered channels and the appropriate reference channels. ‘Dark’ (no signal) and ‘Bright’ (homogeneous, unfiltered light) calibration images were acquired for each day of experiment to account for camera noise and effective light splitting. Single cells are segmented by the Cellpose^36^ library and only the coordinates from segmented cells are used to generated the phasor distribution, while the median phasor position for each cells is displayed as a single point in a separate graph. Images were false colored with the angle (phase) of the calculated phasor position, which linearly correlates with average emission wavelength. As a result, cells display a color that depends on the bioluminescent reporter they express (green for YeNL, cyan for CeNL).

### Fluorescence microscopy imaging and analysis

Cells were seeded in 8-well chambers (Cellvis) at 1 × 10^5^ cells per well the day before imaging. Cells were incubated with organelle-targeting dyes for 3 h before starting collecting data, specifically with TMRM (Tetramethylrhodamine, Methyl Ester, ThermoFisher) at a final concentration of 100 nM and Lysotracker Deep Red (ThermoFisher) at a final concentration of 50 nM. Data were collected with Zeiss LSM880 inverted microscope, using the 32-channel spectral detector module (spectral range 409-690) in Photon-counting mode. The objective used was a Zeiss 63X/1.4 NA Oil objective, pixel size 130 nm, 2.05 μs pixel dwell time, frame size 1024×1024 pixels and to collect the data was used the sum of 8 lines. We excited simultaneously with three laser lines (458 nm, 514 nm, 594 nm). For every condition, we acquired 10 different fields of views and 3 independent replicates of the experiment were collected.

The data were analyzed with a custom Python code. Each image was unmixed in four channels using the spectral phasor unmixing approach, resulting in 4 images (CeNL, YeNL, TMRM and Lysotracker Deep Red). Single cells were segmented using Cellpose^36^ (pretrained ‘cyto2’ network) and the total intensity for each channel was stored in a Pandas Dataframe and exported to Excel. Graphs and statistical analysis (Outliers identification, One-way ANOVA/Kruskal-Wallis test) were performed using Graphpad Prism 5-9 Software.

### Fluorescence Z-stack imaging

Measurements were taken with Zeiss LSM 880 inverted microscope, using the 32-channel spectral detector module (spectral range 409-690 nm) in Photon-counting mode. The objective used was a Zeiss 10X NA 0.45, pixel size 830 nm, pixel dwell time 2.05 μs, frame size 1024×1024 pixels and to collect the data was used the sum of 8 lines. We collected a z-stack dataset up to 310 μm from the bottom of the well in 10 μm increments.

## Results

### BRET reporter cells enabled readout on M1/M2 stimulation

Macrophage polarization reporters were designed with enhanced Nano-lanterns (eNLs) comprising NanoLuc luciferase^37^ as a bioluminescent donor and different fluorescent proteins as acceptors (BRET reporters)^38,39^. Resolving these fusions is often challenging due to their broad, overlapping spectra and incomplete energy transfer. Recently, we showed that spectrally similar bioluminescent probes could be readily distinguished via spectral phasor analysis.^34^ We thus aimed to use this technique for analyzing macrophage polarization reporters. In this scenario, M1- or M2-specific gene upregulation would drive the expression of a corresponding BRET reporter. Different levels of gene expression would then be readily discerned via spectral phasor analysis (**Figure 1**).

**Figure 1.**
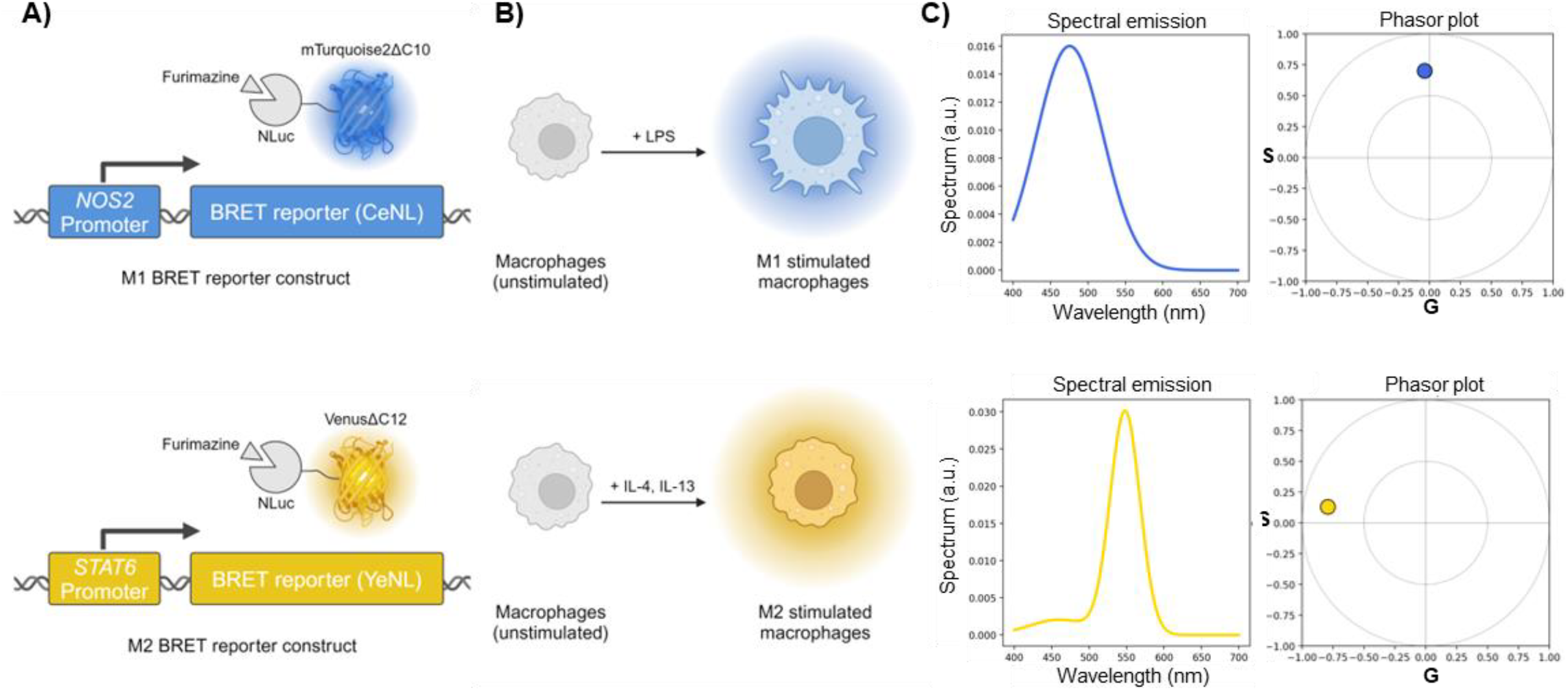
Macrophage polarization reporters together with spectral phasor analysis enable single cell polarization state readout. (A) Cartoon depiction of BRET reporter expression dependent on promoters upregulated in either M1-(NOS2 promoter-CeNL, top) or M2-polarized (STAT6 promoter-YeNL, bottom) macrophages. (B) Cartoon depiction of changes in bioluminescence emission color of cells stably expressing the M1 (top) or M2 (bottom) BRET reporter construct upon stimulation with LPS or IL-4+IL-13, respectively. (C) Emission spectrum (left) and position in the spectral phasor plot (right) of the M1 BRET reporter (CeNL, top) and M2 BRET reporter (YeNL, bottom).

To design the necessary reporters, we focused on combining M1- or M2-specific promoter sequences with different colored BRET probes. The promoters selected were *NOS2* and *STAT6* for M1 and M2 macrophages, respectively (**Figure 1A**). When activated, M1 macrophages produce nitric oxide (NO) to aid in pathogen killing and promoting inflammation. NO production is induced by upregulation of the inducible isoform of nitric oxide synthase (*iNOS* or *NOS2*)^40^. Therefore, *NOS2* gene expression has been used as a reliable marker for identifying M1 macrophage phenotypes^32,40,41^. M2 macrophage polarization relies on the phosphorylation of signal transducer and activator of transcription 6 (STAT6)^42^. While the mechanistic details underlying *STAT6* production and M2 behavior are less well defined, *STAT6* is required for M2 activation by IL-4^43^.

The BRET probes selected were the enhanced Nano-lanterns CeNL and YeNL^39^. These luciferases produce cyan and yellow light, respectively, in the presence of luciferin substrate (furimazine). CeNL and YeNL expression would be driven by the *NOS2* and *STAT6* promoters, respectively (**Figure 1A)**, enabling macrophage polarization to be monitored as shown in the cartoon in **Figure 1B**. We envisioned capturing the emitted light at the single-cell level via spectral phasor analysis, providing a multiplexed readout on macrophage status. Briefly, the emitted light is split in four channels, two of which are filtered using sine- and cosine-shaped emission filters and the other two are unfiltered and used as reference^34^. The resulting images are processed to yield two coordinates, G and S, that encode the emission spectrum in a 2-dimensional space called the spectral phasor space. Spectra with different properties (e.g., average emission wavelength and width) will have distinct phasor locations, as shown in **Figure 1C**. For the desired M1 and M2 reporters, the emission spectrum of *NOS2*-CeNL (**Figure 1C, top left**) would generate a distribution located in the center at the top of the phasor diagram (**Figure 1C, top right**) and the emission spectrum of *STAT6*-YeNL (**Figure 1C, bottom left**) would generate a distribution located on the left side of the phasor plot (**Figure 1C, bottom right**).

To test the overall concept, the engineered reporters were introduced into a model macrophage cell line, RAW264.7 via viral transduction. We then validated BRET reporter expression by inducing M1 or M2 polarization (**Figure 2**). The cell lines were incubated with either M1 (LPS) or M2 (IL-4, IL-13) stimulatory molecules and luminescence was monitored after 24 h via spectral phasor analysis. Our microscopy setup optically encodes the emission spectrum in each pixel to a point of a new space, the phasor space, that is defined by the two coordinates noted above, G and S, as described in **Figure 1**. Converting the spectra into the phasor space allows more facile assignments of complex and highly overlapping spectra. Simultaneous fingerprinting of spectrally similar probes is thus possible. Being a camera-based set up, extended integration times, a feature common to bioluminescent readouts with dim-emitting probes, are also feasible.

**Figure 2.**
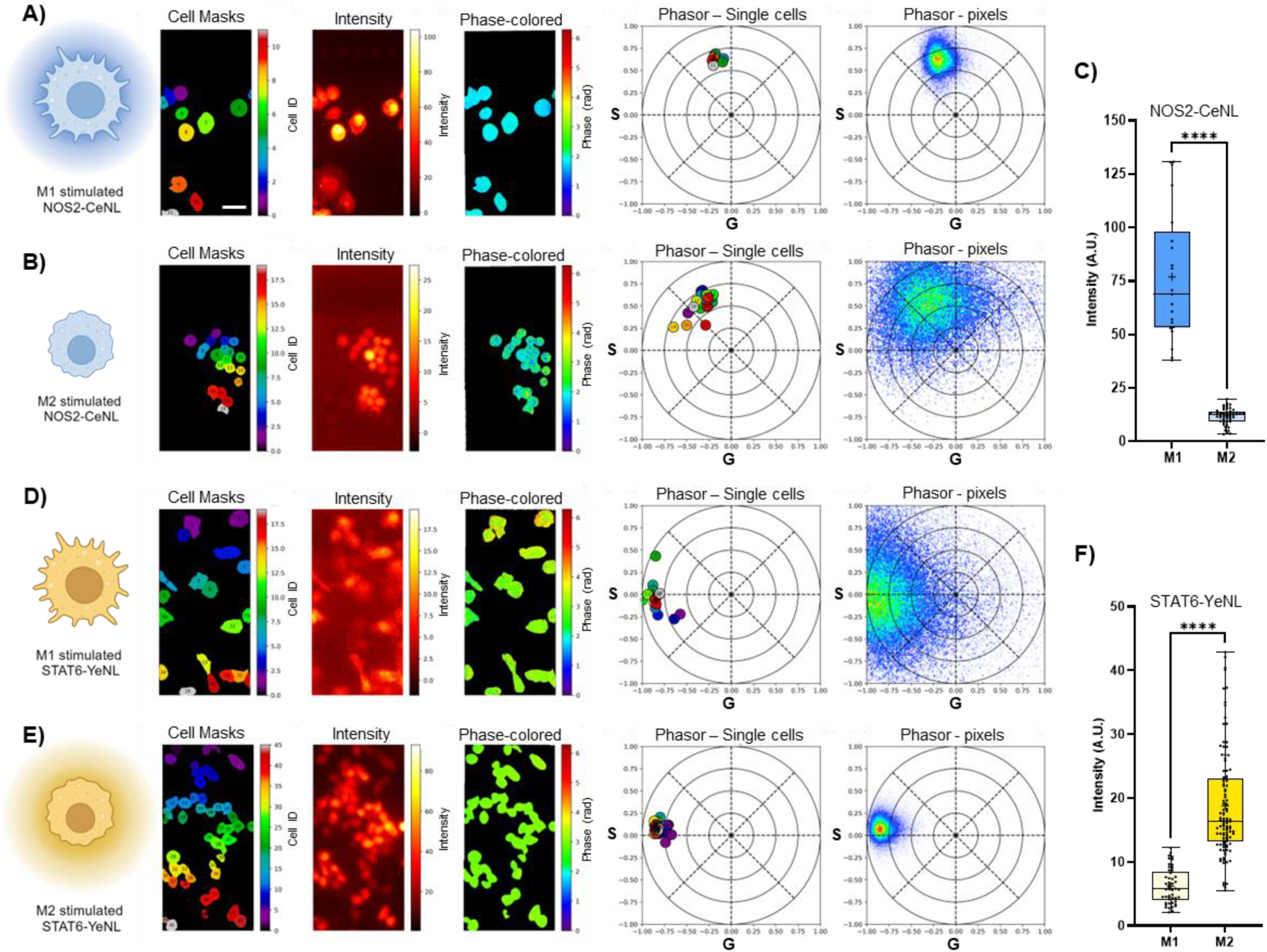
Stimulation with different activators allowed validation of gene expression reporter cell lines via bioluminescence phasors. Cells stably expressing polarization gene expression reporters: NOS2-CeNL (A, B) or STAT6-YeNL (D, E). Cells were stimulated either towards M1 (A, D, IL-4 20 ng/mL and IL-13 20 ng/mL) or M2 (B, E, LPS 5 µg/mL) polarization. For each condition (A, B, D, E) we are reporting, left to right: a cartoon depiction of the cell status upon stimulation, cell segmentation masks, bioluminescence intensity images, false-colored images based on the pixelwise phasor position, phasor plot reporting the median phasor position in single cells and phasor plot of the pixelwise phasor distribution for the whole field of view. Scale bar is 30 μm. Single cell intensity quantification is reported for NOS2-CeNL (C) and STAT6-YeNL (F) cell lines. Data are represented by box and whiskers plots. The box represent the median (solid line) and standard deviation. The whiskers go down to the smallest value and up to the largest. All the points are shown. One-way ANOVA (K-W test), p>0.05 (ns), 0.05>p>0.01 (*), 0.01>p>0.001 (**), 0.001>p>0.0001 (***), p<0.0001 (****). For each condition, a minimum of 2 technical replicates were collected.

For our experiments, each image was constructed by averaging 20 frames at 10 seconds exposure. We then performed single cell segmentation and spectral phasor analysis, in order to calculate the phasor location for every single cell. As expected, the phasor location of the *NOS2*-CeNL cells was located in the top part of the phasor plot, with a more compact distribution when stimulated towards M1 (**Figure 2A, right**), indicative of a larger number of photons collected. A broader distribution was observed (indicative of a smaller number of cells) when the populations was stimulated towards M2 (**Figure 2B, right**). For the *NOS2*-CeNL reporter cells, the increase in bioluminescent output was ∼6.5 fold when stimulated towards M1 compared to when stimulated towards M2, as shown in the boxplots in **Figure 2C**. The opposite trend was observed for the *STAT6*-YeNL cells. A broader distribution in the left side of the phasor plot (**Figure 2D, right**) was observed when the cells were stimulated towards M1, while a more compact distribution resulted when cells were stimulated towards M2 (**Figure 2E, right**). The overall increase in bioluminescent signal was ∼3-fold when this population of cells was stimulated towards M2 compared to M1 (**Figure 2F**). The overall fold change was moderate, but in agreement with similar reports^44^.

Reporter cell line sensitivities to stimulation were also confirmed with other assays. Bulk luminometer measurements showcased successful eNL expression and light emission (**Figure S1**). Upregulation of M1- and M2-specific reporters was further confirmed by direct laser excitation of the fluorescent protein acceptors comprising each BRET construct. These experiments were conducted with cells expressing either *NOS2*-CeNL or *STAT6*-YeNL, along with a 1:1 mixture of the two reporters (**Figure S2A, B**). Additionally, macrophage phenotypes were validated using fluorescent dyes that report on mitochondrial membrane potential and lysosome activity (**Figure S2C, D**). Similar to previous reports, we observed a higher mitochondrial membrane potential^45^ as well as enlarged and more active lysosomes^46^ in M1-stimulated macrophages compared to unstimulated or M2-stimulated cells. These results establish the BRET reporter lines and phasor analysis as a robust imaging platform for monitoring macrophage phenotypes.

### Bioluminescent phasors enable multiplexed imaging of reporter cell populations

Since spectral phasor analysis enables single cell readouts on BRET expression, we next examined whether the two reporters (*NOS2*-CeNL and *STAT6*-YeNL) could be distinguished and quantified in a single sample. We co-cultured *NOS2*-CeNL and *STAT6*-YeNL cells and exposed them to M1 (**Figure 3A-C**) or M2 (**Figure 3D-F**) stimulation, classifying their status based on phasor location. High intensity readouts for M1-stimulated macrophages were observed when the co-cultures were incubated with LPS. Cell segmentation was performed to obtain the average phasor position for individual cells following LPS treatment (**Figure 3C**). Clusters of phasor positions were observed that corresponded to the emission phasor position for NOS2-CeNL (**Figure 3C**, first phasor plot). Pixel-wise phasor distributions are also shown (**Figure 3C**, second phasor plot). As shown in **Figure 3C**, signal was also observed from the *STAT6*-YeNL reporter cells, as the intensity values for M2 macrophages can reach up to 30% of the total signal (**Figure 3A, right**).

**Figure 3.**
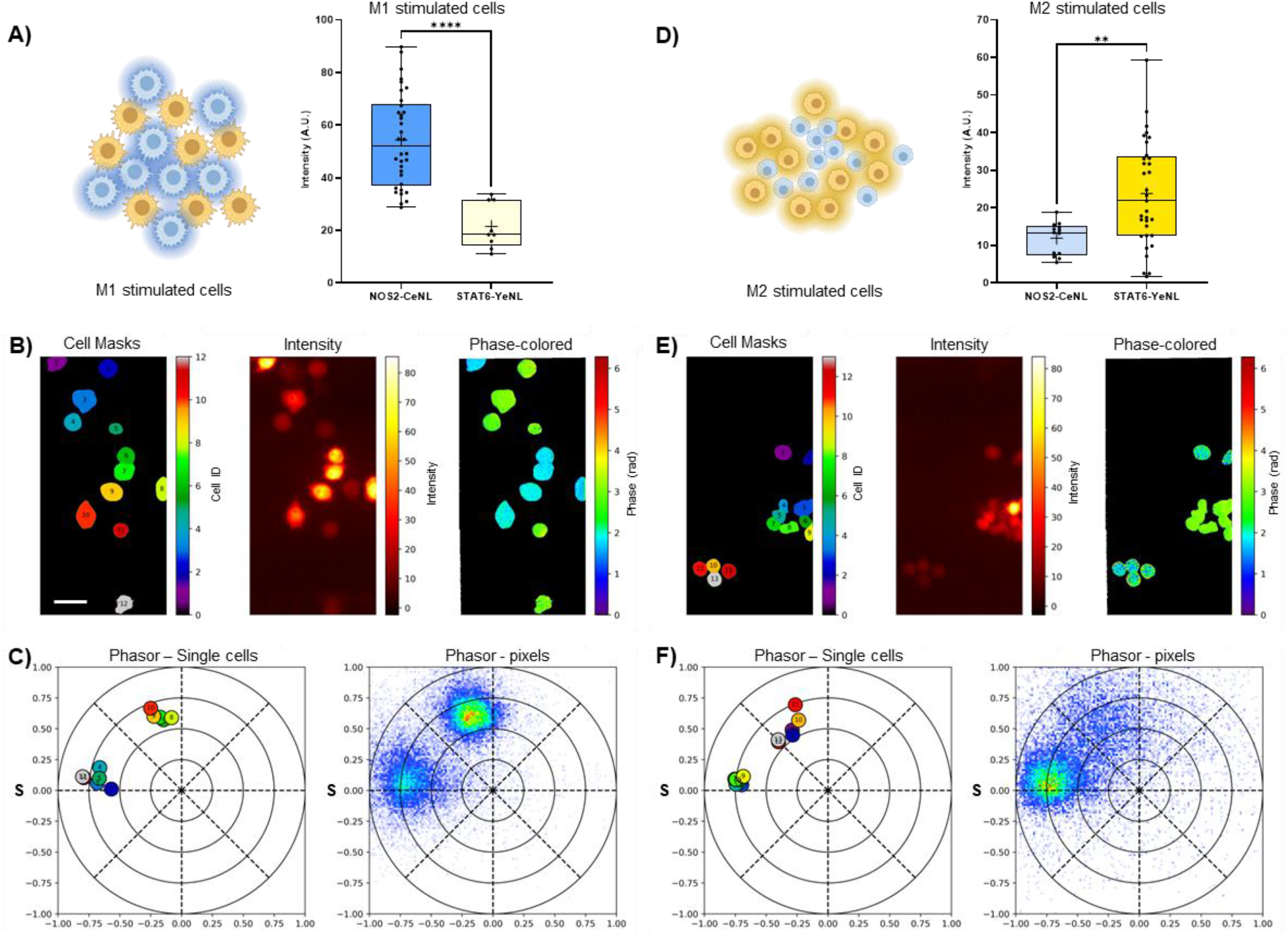
Bioluminescence spectral phasor approach can distinguish polarization of a mixed population of reporter cells. Cartoon depiction of a mixed cell population stimulated towards M1 (A, left) or M2 (D, left) polarization and quantification of the bioluminescence intensity for the two cell reporters (A, D, right). Data are represented by box and whiskers plots. The box represent the median (solid line) and standard deviation. The whiskers go down to the smallest value and up to the largest. A ll the points are shown. One-way ANOVA (K-W test), p>0.05 (ns), 0.05>p>0.01 (*), 0.01>p>0.001 (**), 0.001>p>0.0001 (***), p<0.0001 (****). (B, E) Left to right, masks of the segmented cells, bioluminescence intensity and phase-colored images of a mixed cell population stimulated towards M1 (B) or M2 (E). (C, F) Average phasor location of single cells (left) and of the whole field of view (right) of a mixed cell population stimulated towards M1 (C) or M2 (F). Scale bar is 30 μm. For each condition, a minimum of 5 technical replicates were collected.

Similar experiments were performed with M2-stimulating cytokines. As shown in **Figures 3D**, large photon outputs were observed when co-cultured cells were treated with IL-4/IL-13. A more compact distribution was observed for *STAT6*-YeNL phasor position. The contribution from the other reporter (*NOS2*-CeNL), in this case, was less prevalent. While both cell lines undergo polarization (also confirmed by cell morphology), only the cells expressing the appropriate gene expression reporter display an increase in intensity, as clearly shown in the boxplots in **Figures 3A,D**.

### Macrophage polarization can be monitored in 3D tissue mimic

Finally, we investigated the applicability of the imaging method in a 3D structure to mimic tissue organization. An important feature of our approach is that the bioluminescent probes and spectral phasor analysis are compatible with serial, long-term imaging in heterogeneous environments. To demonstrate feasibility, we embedded mixtures of *NOS2*-CeNL and *STAT6*-YeNL reporter macrophages in a collagen matrix at varying depths. M1 and M2 stimulation treatments were dispensed dropwise at the top of the collagen matrix, generating a top-to-bottom stimulation gradient, resembling *in vivo* conditions (**Figure 4A**).

**Figure 4.**
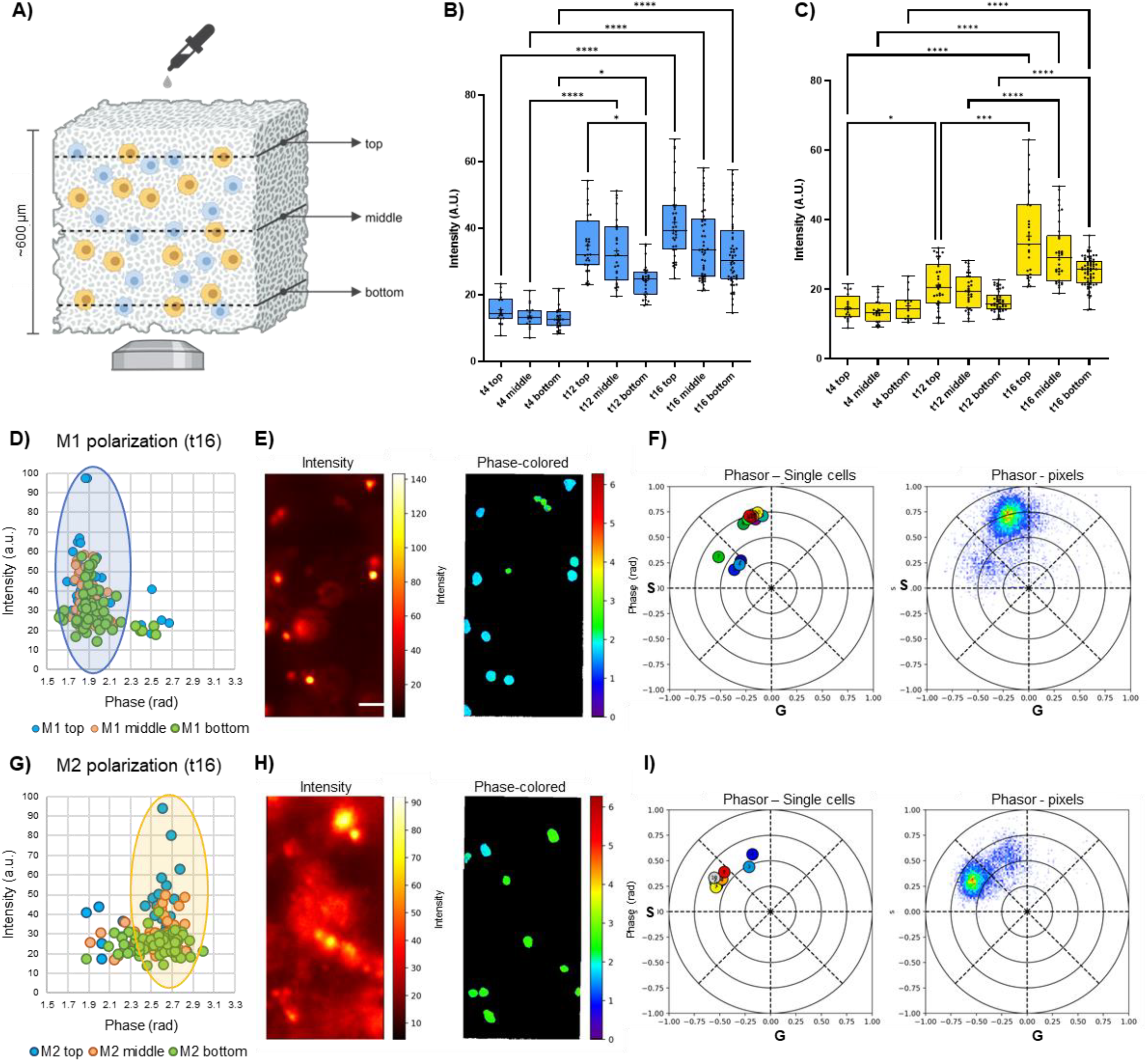
Imaging macrophage reporter cells in 3D tissue mimic. (A) Cartoon depiction of a mixed cell population embedded in a 3D collagen matrix. Luciferase substate (furimazine) was added from the top and detection is collected from the bottom of the sample. Measurements were taken at 3 different depths (top, middle, bottom) as described in the Methods section. Quantification of bioluminescence intensity of the NOS2-CeNL reporter when stimulated towards M1 (B) and of the STAT6-YeNL reporter when stimulated towards M2 (C). Measurements were taken at different time points after incubation with cytokines (4, 12 and 16 hours) and at different depths. Data are represented by box and whiskers plots. The box represent the median (solid line) and standard deviation. The whiskers go down to the smallest value and up to the largest. All the points are shown. One-way ANOVA (K-W test), p>0.05 (ns), 0.05>p>0.01 (*), 0.01>p>0.001 (**), 0.001>p>0.0001 (***), p<0.0001 (****). Scatter plots of the bioluminescence intensity as a function of the spectral phase (in radians) of the cells imaged after 16 hours of cytokine incubation for M1 (D) and M2 (G) stimulation. M1-polarized NOS2-CeNL cell population is highlighted by the blue oval in (D), and M2-polarized STAT6-YeNL cell population is highlighted by the yellow oval in (G). Bioluminescence intensity (left) and phase-colored images (right) of the “top” depth of a mixed cell population after 16 hours of stimulation towards M1 (E) or M2 (H). Scale bar is 60 μm. Average phasor location of single cells (left) and of the whole field of view (right) of the mixed cell population stimulated towards M1 (F) or M2 (I), corresponding to the field of views shown in E and H respectively. For each condition, a minimum of 2 biological replicates (each with 2 technical replicates) were collected.

Images were acquired 4, 12, and 16 h post-cytokine addition at several planes. Intensity distributions show an increase in NOS2 (**Figure 4B**) and STAT6 expression (**Figure 4C)**, as a function of time for M1 and M2 stimulation, respectively, with a significant increase at 12 h for NOS2 and at 16 h for STAT6.

Furthermore, in particular at the 12-hour and 16-hour time points, a gene expression gradient can be appreciated from top to bottom as a result of the top-to-bottom localized application of the stimulant. Sixteen hours after cytokine addition, scatter plots of the spectral phase averaged from the phasor plot (which correspond to single cells show selective activation of either *NOS2*-CeNL (**Figure 4D**) or *STAT6*-YeNL (**Figure 4G**) following M1 or M2 activation, respectively. Scatter plots related to the 4 h (t4) and 12 h (t12) time points were also calculated (**Figure S3**). Additionally, we excited directly the fluorescent acceptors of the BRET reporters to confirm the presence of both cell lines. As shown in **Figure S4** we collected z-stack datasets of the spectral emission upon different stimulations, 3 hours and 10 hours after addition of the stimuli.

## Discussion

Methods to visualize immune function must capture a spectrum of behaviors in physiologically relevant environments. This is no easy task, considering the dearth of methods suitable for continuous and, ideally, noninvasive recording. We sought to fill this void using bioluminescent technologies. Bioluminescence is well suited for real-time monitoring, but few tools can report on the complexities of immune function, maintaining single-cell resolution.

We have demonstrated a novel approach to monitoring macrophage polarization using genetically engineered BRET expression reporters and spectral phasor analysis. The BRET reporters enabled a multiplexed readout on macrophage polarization in cell mixtures and over time. This strategy can also be used to monitor macrophage polarization in a 3D tissue mimic, leading the way to applications *in vivo*. Additionally, the polarization BRET reporters can be combined with fluorescence imaging via direct excitation of the reporters themselves or with exogenous fluorescent dyes. Such experiments provide a multi-dimensional readout on macrophage polarization.

While spectral phasor analysis is useful for monitoring BRET reporter expression in polarized macrophages, additional optimization is warranted. Future studies would benefit from singular constructs and additional reporters to provide a more comprehensive readout on macrophage phenotype. Such a strategy also requires methods to stably integrate multiple reporter genes per cell to enable single cell analysis of a variety of biomarkers.

In conclusion, bioluminescent enhanced Nano-lantern reporters combined with spectral phasor imaging can be used to monitor macrophage polarization dynamics in complex samples. This work provides a blueprint for single-cell bioluminescent readouts of multiplexed gene expression.

## Acknowledgments

Cartoon panels were created with BioRender.com. This work was supported by the U.S. National Institutes of Health (R01 GM107630 to J.A.P.) and the Paul G. Allen Frontiers Group (to J.A.P, M.A.D., L.S, G.T.). M.X.N. was supported by a NSF Graduate Research Fellowship (DGE-1321846). We thank Prof. Enrico Gratton and members of the Laboratory of Fluorescence Dynamics (LFD, UCI) for helpful discussion.

## Author contributions

MAD and JAP conceived the project idea. MXN and TKS generated the reporter constructs. MXN and GT prepared the biological samples. MXN and GT performed the experiments. LS wrote the codes to analyze the data and optimized the imaging set up. MXN and GT analyzed the data. All authors contributed to the writing of the manuscript. All authors have given approval to the final version of the manuscript.

