## Supplementary Figures for "Monitoring macrophage polarization with gene expression reporters and bioluminescence phasor analysis"

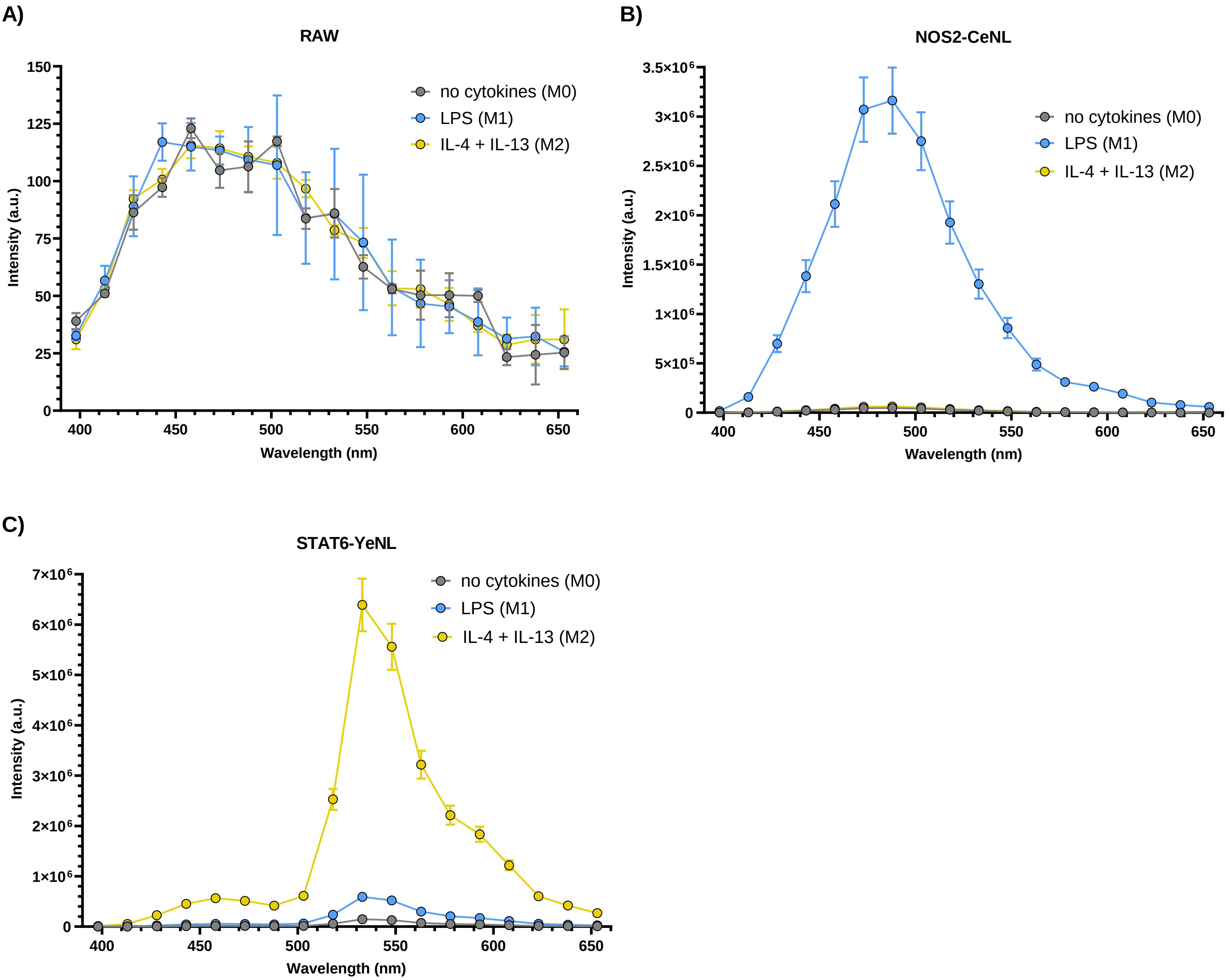


**Supplementary Figure 1.** Untransduced RAW macrophages **(A)**, *NOS2*-CeNL stable RAW cell line **(B)** and *STAT6*-YeNL stable RAW cell line **(C)** were incubated with furimarize and their luminescence was measured with a luminometer in absence of stimuli (grey), in the presence of LPS (blue) to stimulate towards M1 or in the presence of IL-4 + IL-13 (yellow) to stimulate towards M2. Data are represented as mean and error (standard deviation).

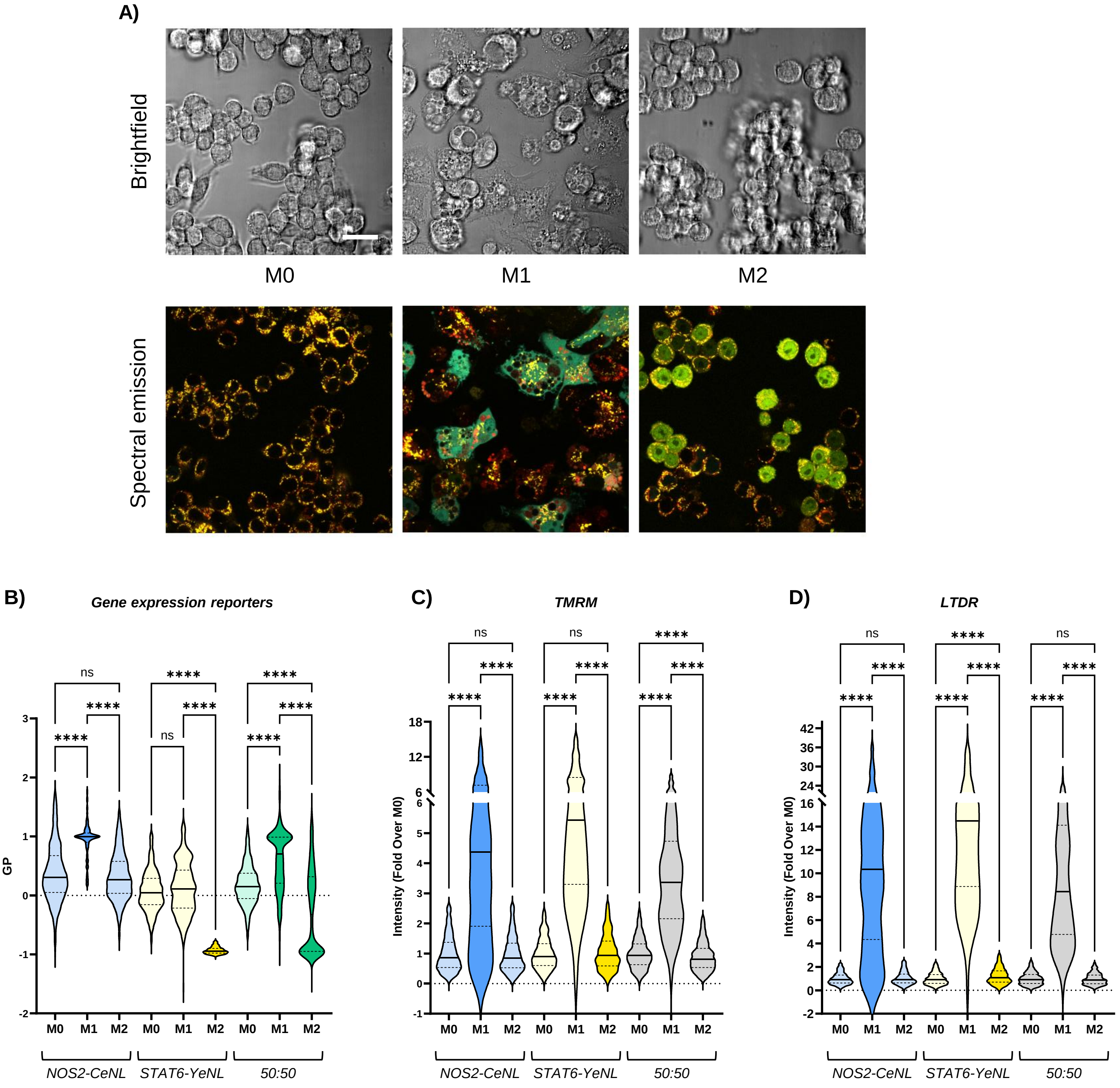


**Supplementary Figure 2. (A)** Brightfield (top row) and spectral emission (bottom row) of a 50:50 mixture of *NOS2*-CeNL and *STAT6*-YeNL macrophages labeled with organelle-targeting dyes (TMRM, Lysotracker Deep Red) under different stimulation conditions (M0=no stimulation, M1=stimulation with LPS, M2=stimulation with IL4-IL13). Scale bar is 20 µm. **(B)** Generalized Polarization (GP) of the intensities emitted by the reporters upon direct excitation of the fluorophores (mTurquoise2ΔC10 for *NOS2*-CeNL and VenusΔC12 for *STAT6*-YeNL) under different stimulation conditions (M0, M1, M2). GP=1 represents 100% *NOS2*-CeNL expression, GP=-1 represents 100% *STAT6*-YeNL expression. **(C)** Intensity of the mitochondria probe (TMRM, TetraMethylRhodamine Methylester. Higher intensity=higher mitochondrial membrane potential) across the different macrophages lines, under different stimulations (M0, M1, M2). **(D)** Intensity of the lysosome probe (LTDR, LysoTracker Deep Red. Higher intensity=lower lysosomal pH) across the different macrophages lines, under different stimulations (M0, M1, M2). Data are represented by violin plots. The solid line represent the median and the dotted lines represent the quartiles. One-way ANOVA (K-W test), p>0.05 (ns), 0.05>p>0.01 (*), 0.01>p>0.001 (**), 0.001>p>0.0001 (***), p<0.0001 (****).

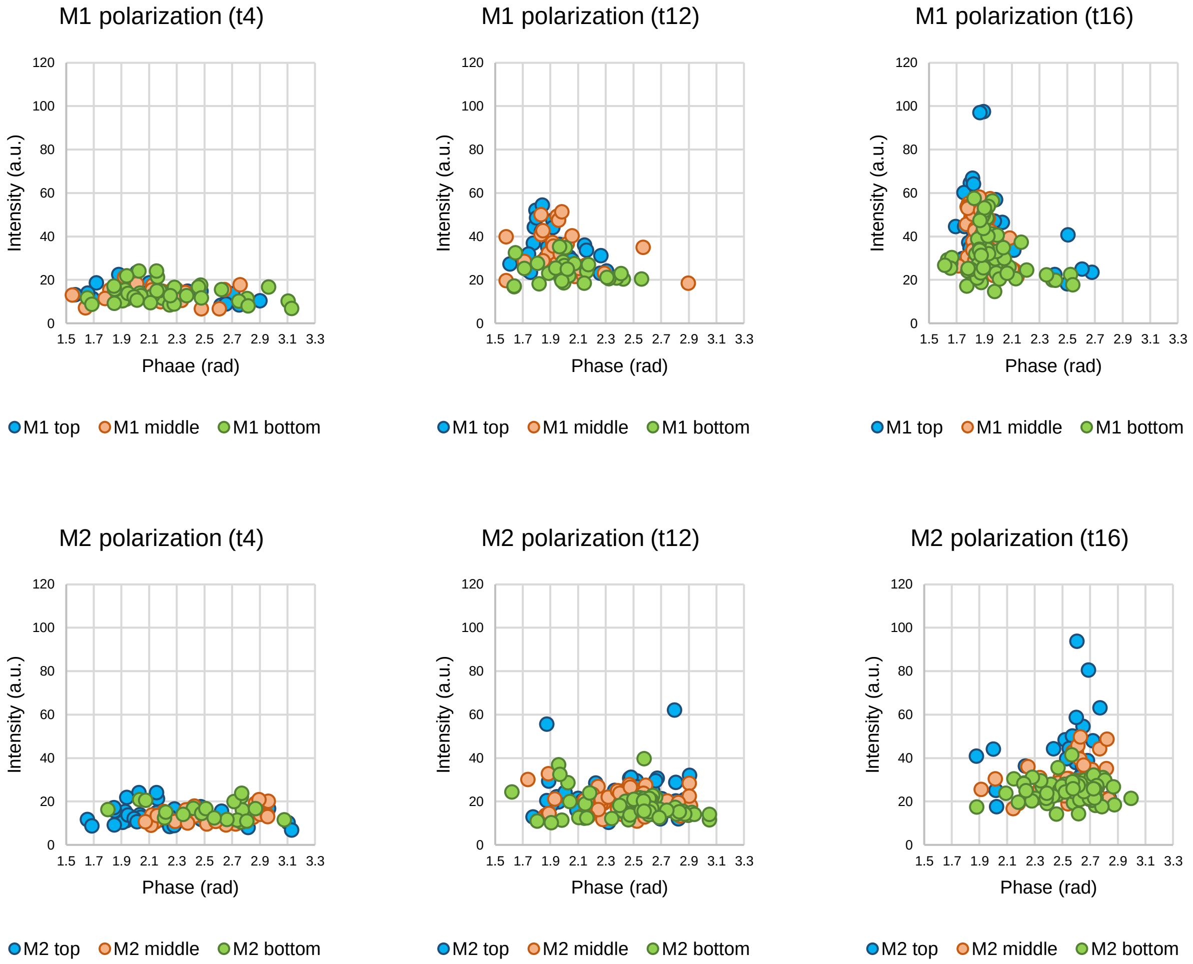


**Supplementary Figure 3.** Scatter plots of the bioluminescence intensity as a function of the spectral phase (in radians) of the cells imaged after 4 (t4), 12 (t12) and 16 (t16) hours of cytokines incubation for M1 and M2 stimulation. Each dot represents a single cell.

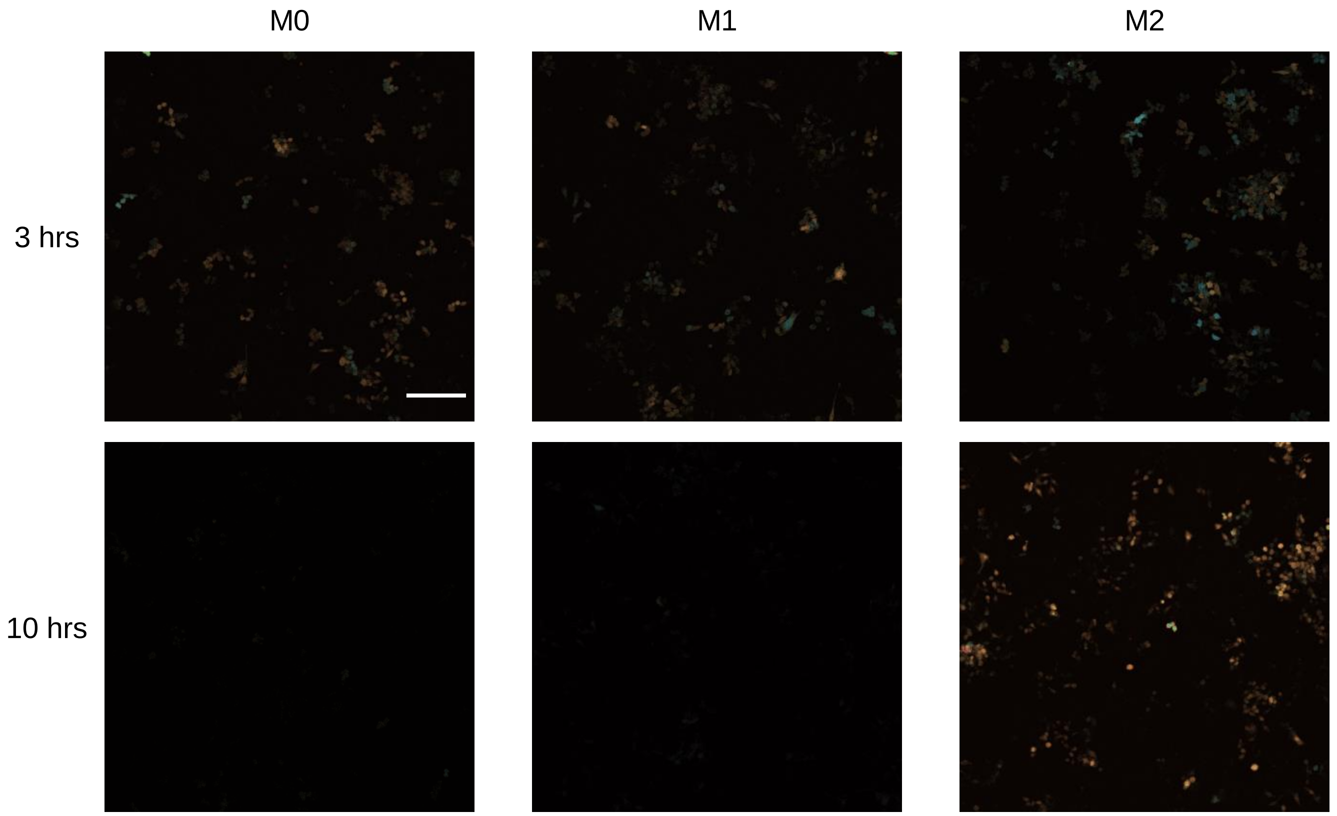


**Supplementary Figure 4.** Spectral emission of a mixed population of *NOS2*-CeNL and *STAT6*-YeNL embedded in collagen. **Top row**: z stack of the spectral emission after 3 hours of M1, M2 or no stimulation (M0). **Bottom row**: z stack of the spectral emission after 10 hours of M1, M2 or no stimulation (M0). Scale bar is 130 μm.
